# Quality of life of people seeking for acupuncture treatment

**DOI:** 10.1101/2020.07.10.196907

**Authors:** Reginaldo Silva-Filho, Marcia Kiyomi Koike, Gizelda Monteiro da Silva

## Abstract

**Background:** The use of Acupuncture, part of Chinese Medicine, has grown as well as the need to understand its effects. The Quality of Life (QOL) assessment is an important way to have a global view of the patient.

**Objective:** To assess the QoL of people who seek acupuncture at the outpatient clinic of an acupuncture school.

**Methods:** QOL assessment using WHOQOL-BREF applied only once to people who voluntarily sought for acupuncture at the general outpatient clinic of Faculdade EBRAMEC in 2016.

**Results:** People who sought for acupuncture treatment presented lower mean QoL value in all domains compared to normative values for Brazil, with physical domain presenting lower values than the other domains. It was also observed that male patients had higher values in the physical, psychological domains and general QOL compared to women.

**Conclusion:** Our data demonstrated that people seeking for acupuncture treatment presented lower values of QOL in comparison to a referential study with a Brazilian population, providing for the first time reference values measured by WHOQOL-bref, for patients seeking for acupuncture in the city of São Paulo.

## 1. Introduction

Chinese Medicine, a medical rationale [1] that has a history of more than 5000 years of practice, presents a peculiar way of understanding physiology and pathology of the human being [2], frequently through observations and connections related to external environment and the body, but also related to the connections of different parts of the body with each other and as a whole, concept that is described in the West as a holistic approach [3,4] As for the treatments, Chinese Medicine offers different approaches: Acupuncture and Moxibustion; Chinese herbal medicine; Chinese diet therapy; Chinese massage therapy (Tui Na); Chinese Body Arts (Qi Gong and Tai ji Quan) [5].

Acupuncture has been part of the National Policy of “Integrative and Complementary Practices in Health” (PICS) since 2006 [6] and its use through Brazilian Unified Health System (SUS) has grown in the official data, presenting itself as one of the most used treatments among the PICS, second only to phytotherapy. Researchers identified that the use of acupuncture in SUS is still small compared to its use in the private health system [7].

The concept of “Quality of life” has been in use for the evaluation of therapeutic methods [8-12], including those identified as complementary, alternative or integrative [13-18], where acupuncture is part. World Health Organization (WHO) Quality of Life group defines it as “individuals’ perception of their position in life in the context of the culture and value systems in which they live and in relation to their goals, expectations, standards and concerns”[19]. This wider approach allows an improvement on the communication between patients and health professionals during the evaluation process, which leads to a gain in the identification and establishment of goals and priorities for the treatment [20-23].

There are different instruments translated, adapted and validated in Portuguese, including those developed and recommended by the WHO, such as World Health Organization Quality of Life Questionnaire 100 (WHOQOL-100), translated into at least 20 languages [24], and the World Health Organization Quality of Life Questionnaire - Bref (WHOQOL-BREF), a more practical and direct version, with the same characteristics [25,26].

Specifically to acupuncture, the use of “Quality of Life” assessment instruments has been frequent to compare its effects, however there is no reference or normative value for those who seek acupuncture treatment. In this sense, the present study aims to assess the quality of life of people seeking for acupuncture treatment in an outpatient clinic of an acupuncture school, in addition to presenting a description of the age and gender profile of people who seek for acupuncture treatment, and compare the quality of life indexes of these people with the normative data indexes of a Brazilian population.

## 2. Materials and Methods

This study was carried out through an observational, descriptive, transversal research with a quantitative approach, through WHOQOL-BREF instrument data evaluation after being applied to those who voluntarily sought for acupuncture treatment at the general outpatient clinic of the EBRAMEC College (Brazilian School of Chinese Medicine) and answered the instrument in a single moment - directly before the first appointment, during 2016.

WHOQOL-BREF [24], the selected instrument for this study, contains 4 domains and 26 questions, separated into physical, psychological, social, environmental, which application presented satisfactory psychometric properties of internal consistency, discriminant validity, criterion validity, concurrent validity and test-retest reliability.

Data collection started after being approved by the Research Ethics Committee of - Instituto de Assistência Médica do Servidor Público Estadual - IAMSPE, obtaining the Certificate of Presentation for Ethical Appreciation (CAAE): 61575916.1.0000.5463. Following all the guidelines and regulatory standards present in the Resolution of the National Health Council No. 466/2012.

Data were properly tabulated, evaluated and analyzed according to the recommendations of Brazilian WHOQOL Group, responsible for the translation, cultural adaptation and validation of WHOQOL-BREF in Portuguese language. Participants were divided into age groups according to the normative reference for the Brazilian population [27]: <20 years, ≥ 20 years and <30 years, ≥ 30 years and <45 years, ≥ 45 years and <65 years and ≥ 65 years.

Data obtained in the present study (Acupuncture Group) were also compared with the normative indexes of a Brazilian population using WHOQOL-BREF, according to the Brazilian WHOQOL Group (Brazil Group) [27] and with the basic data of an older adults population (Porto Alegre Group) [28]. The surveyed variables were analyzed according to the recommendations of Brazilian WHOQOL Group [29].

For the Acupuncture Group, there were some criteria:

### 2.1 Inclusion Criteria

- Patients of the general acupuncture clinic of Faculdade EBRAMEC;
- Being treated by acupuncture at least once on the previous mentioned clinic after answering the evaluation instrument.

### 2.2 Exclusion Criteria

- Teachers, employees, and students of Faculdade EBRAMEC.

### 2.3 Statistical Analysis

The results were presented as mean and standard deviation for the continuous variables and, frequency and proportion for categorical variables. Statistical analysis was performed using the SPSS 22.0 software.

For the proofing of existence of dependency relationships between categorical variables, the chi-square test was used. The t-Student or Mann-Whitney tests were used to compare continuous variables between two independent groups, as in the case of comparisons between males and females.

Wilcoxon test was used for paired variables, such as the comparison between results in pairs of WHOQOL-BREF domains. Kruskal-Wallis test was used to compare three or more independent groups, such as in different age groups and, in the presence of significant differences, Bonferroni correction or procedure was used.

Spearman test was used to verify the correlation between the variables related to the results of each of the domains of the questionnaire. All variables were tested to find out whether they had a normal distribution by Kolmogorov-Smirnov test, in order to decide which of the tests described above would be used. In the present study, a level of statistical significance of less than or equal to 5% was adopted.

## 3. Results

During the period of study, a total of 353 patients who sought treatment by acupuncture at the general outpatient clinic of Faculdade EBRAMEC were analyzed, including 111 (31.4%) male and 242 (68.6%) female. The proportion of female patients was higher than that of male patients (chi-square adherence test with p <0.001).

Among the 353 patients, the mean age was 52.9 (± 17.8) with a very wide age variation with a minimum of 7 and a maximum of 90 years of age, as shown in Table 1.

**Table 1.** Data on age group of participants.

| Age Group | N | % |
| --- | --- | --- |
| < 20 years | 12 | 3.40 |
| $\geq 20$ years e < 30 years | 26 | 7.37 |
| $\geq 30$ years e < 45 years | 73 | 20.68 |
| $\geq 45$ years e < 65 years | 141 | 39.94 |
| $\geq 65$ years | 101 | 28.61 |
| Total | 353 | 100 |

The indexes of general Quality of Life and each of the domains of WHOQOL-BREF, for the 353 individuals are shown in Table 2.

**Table 2.** Indexes of General Quality of Life and each of the domains of patients seeking for acupuncture treatment.

| WHOQOL<br>Index | Domain |
| --- | --- |
| General Quality of Life | 59 (13,5) |
| Physical Domain | 54,1 (18,2) |
| Psychological Domain | 61,5 (17,2) |
| Social Domain | 62,0 (18,6) |
| Environment Domain | 58,3 (14,1) |

Mean (standard deviation)

The results of each WHOQOL-BREF domain and the general Quality of Life in relation to male and female patients are shown in Table 3. Higher values were observed in males for physical and psychological domains and quality of life compared to female patients.

**Table 3.** Comparison of the values of each domain and quality of life in relation to gender.

| WHOQOL-BREF | Male | Female | P value |
| --- | --- | --- | --- |
| Physical | 56.7 (16.8) | 52.9 (18.7) | 0.042 |
| Psychological | 66.9 (16.5) | 59.0 (17.0) | < 0.001 |
| Social | 63.7 (20.9) | 61.3 (17.5) | 0.084 |
| Environment | 59.7 (13.4) | 57.7 (14.4) | 0.186 |
| Quality of Life | 61.8 (13.6) | 57.7 (13.2) | 0.004 |

Statistical test: Mann-Whitney

Regarding the different age groups, it was only observed a difference in relation to the Psychological Domain. Age group ≥ 30 years and <45 years showed lower values than age group ≥ 45 years and <65 years (p <0.001) and less than age group ≥ 65 years (p = 0.030).

Physical Domain values were lower than all other domains (with p <0.001 for all comparisons). No differences were identified between the values of Psychological and Social Domains (p = 0.494). And, Environment Domain presented lower values than Psychological and Social Domains (with p <0.001 for each of the two comparisons).

All relationships between the domains showed positive correlations, indicating that patients with higher values in one of the domains also present higher values in the other domains and in the Quality of Life (Table 4).

**Table 4.** Correlation between Domains and Quality of life of participants.

| WHOQOL-BREF | Psychological | Social | Environment | Quality of Life |
| --- | --- | --- | --- | --- |
| Physical | 0.567 | 0.351 | 0.477 | 0.759 |
| Psychological |  | 0.562 | 0.573 | 0.857 |
| Social |  |  | 0.428 | 0.750 |
| Environment |  |  |  | 0.752 |

Statistical test: Spearman correlation

### 3.1 Comparison with normative data for Brazil

For comparison with normative values for the Brazilian population [27], an adjustment was made to analyze only patients with similar age groups, totaling 240 (68% of patients), considering ages between 20 to 64 years. In this analysis, patients who sought for acupuncture treatment (Acupuncture Group) had a significantly lower mean value in all Domains, except for the analysis of the Environmental Domain, compared to the normative sample (Group Brazil) (Table 5).

**Table 5.** Comparison of each domain values between reference of the Brazilian population and people seeking for acupuncture treatment.

| WHOQOL-BREF | Brazil | Acupuncture | P value |
| --- | --- | --- | --- |
| Physical | 58.9 (10.5) | 55.5 (18.3) | 0.007 |
| Psychological | 65.9 (10.8) | 61.0 (17.1) | < 0.001 |
| Social | 76.2 (18.8) | 61.5 (19.2) | < 0.001 |
| Environment | 59.9 (14.9) | 58.1 (14.5) | 0.097 |

Statistical test: Mann-Whitney

For male patients, only the Social Domain of those who sought for acupuncture treatment had a lower mean value than the normative data of Brazilian population (p = 0.002). For female patients, the mean value of all domains, except for the Environment Domain, was lower than data of Brazilian population.

Regarding the different age groups, it was possible to identify that for age group of 20-29 years, the Psychological and Social Domains presented lower mean values; for age group of 30-44 years, only the Physical Domain did not present a lower mean value; and for age group 45-64, only the Social Domain had a lower mean value compared to the normative values [27] (Figure 1).

**Figure 1.**
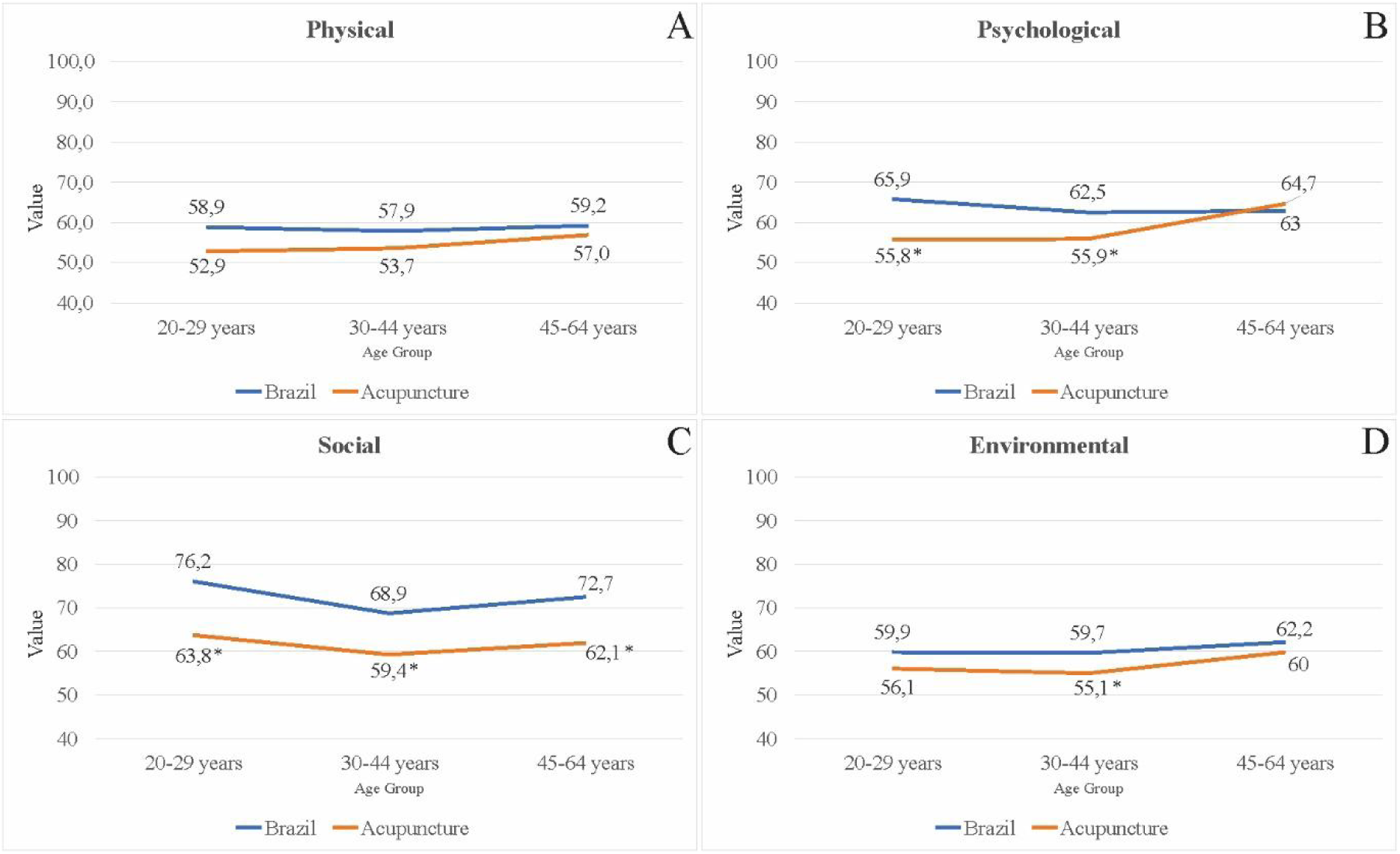
Curves of the WHOQOL-BREF domains, according to age group, in Brazil and Acupuncture Groups. A-Physical; B-Psychological; C-Social; D-Environmental.

This Figure depicts all domains and in all age groups analyzed, the Quality of Life values of the normative data of the Brazilian population were higher than those presented by patients in search of acupuncture, with the exception of data for the psychological domain in the age group between 45 and 64 years old.

### 3.2 Comparison with data from an older population in Porto Alegre

For comparison with an elderly population, it was used data from patients over 60 years old from the Porto Alegre region. 130 (36.83% of the total patients) were selected from the Acupuncture group, adjusting the analysis by age group. It was possible to observe that in all domains, the mean values of individuals who sought care for Acupuncture were lower than the means of individuals in the Porto Alegre region [28] (Table 6).

**Table 6.** Comparison of the domains values of people who sought for acupuncture treatment and reference values for elderly people.

| WHOQOL-BREF | Porto Alegre | Acupuncture | P value |
| --- | --- | --- | --- |
| Physical | 73.84 (14.60) | 52.86 (17.02) | < 0.001 |
| Psychological | 73.71 (12.72) | 64.87 (15.09) | < 0.001 |
| Social | 73.46 (14.85) | 63.78 (16.43) | < 0.001 |
| Environment | 71.94 (12.60) | 58.68 (13.02) | < 0.001 |

Statistical test: Mann-Whitney

## 4. Discussion

The present study offers reference values for the Quality of Life of people seeking for acupuncture treatment based on the population of São Paulo, southeastern Brazil, indicating that the Quality of Life indexes and the respective domains, with the exception of environmental domain, were lower compared to the normative data of the population of Brazil.

The largest age group with patients who sought for acupuncture treatment was between 45 and 65 years old (40%), which is very similar to a study carried out in Santa Catarina [30], where most patients (65%) were from age group of 45 years and older. It is also interesting to point that this is exactly the age group which presented a quality of life value, specifically in the psychological domain, higher in the Acupuncture Group in relation to the normative data for the Brazilian population. This data can be analyzed by the increasing incidence of common mental disorders in older patients [31,32].

According to the principles of Chinese Medicine, the signs and symptoms presented by older patients tend to be related to the concept of Deficiency Syndromes [33], which could be related to the fact that in the comparison for people over 60, data from people seeking for acupuncture treatment were lower in all WHOQOL-BREF domains.

Our study identified that male patients had superior general quality of life indexes including the respective domains in relation to female patients, as also observed in other previous studies from different countries [34,35].

Physical domain presented significantly lower values than all other domains, which is in line with the data presented in China [36,37] and Brazil [38], which presented 58% and 60%, respectively, of the total acupuncture patients analyzed with musculoskeletal complaints, which reflects on a lower physical domain quality of life, as also verified in other studies [39-41]

Acupuncture, as part of Chinese Medicine, has thousands of years of practice, however scientific studies are much more recent, starting in the 18th century with some simple studies, growing into seeking to evaluate the real effectiveness of acupuncture and its physiological and underlying biological factors [42].

Researchers indicate that clinical studies with Acupuncture still need to improve so that the results of basic research can actually be transformed into clinical results [43]. It can be considered that much of this situation is due to the fact that nowadays, what is considered the “gold standard” for evaluating the clinical features of treatments are double blinded, randomized, controlled trials.

Researchers from Chengdu [44], China, describe that in relation to the assessment of clinical results from Acupuncture, research should focus on more reliable measurements, such as symptom and function scales, in addition to assessing quality of life.

As highlighted by Scognamillo-Szabo (2011) [45], Acupuncture theory suggests that health is dependent on psycho-neuro-endocrine functions, under the influence of the genetic code and extrinsic factors such as nutrition, lifestyle, climate, quality of the environment, among others, which also can be seen as understanding the effects of Acupuncture through the use of appropriate instruments for the assessment of Quality of Life, which contemplates different aspects of the human being.

One of the limitations of the study may be in relation to the disproportionate sample in relation to the gender with 111 (31.4%) male and 242 (68.6%) female, which can be understood by historical-cultural issues where men are seen as invulnerable and end up not seeking proper health care, as with women [46,47].

The data in this study provide a new view on people who seek for acupuncture treatments and allow future studies to have a reference for comparison in relation to different groups of people who seek different therapeutic approaches, such as complementary, alternative or integrative practices, as a reference value.

## 5. Conclusions

Data from the present study demonstrated that the mean quality of life of people seeking for acupuncture treatment is lower than that presented in a reference study with a Brazilian population. Thus, this study provides, for the first time, quality of life reference values, measured by WHOQOL-bref instrument, for patients seeking for acupuncture treatment at an institution in the city of São Paulo, southeastern Brazil.

These data can be used as a reference for future studies through comparison with patients seeking for other therapeutic approaches, Western or Eastern, or even as a possible normative reference value for research comparing pre and post therapeutic intervention.

## Acknowledgments

We would like to acknowledge the support from staff of Faculdade EBRAMEC on providing the raw data from the patients.

